# A Neuromuscular Model of Human Locomotion Combines Spinal Reflex Circuits with Voluntary Movements

**DOI:** 10.1101/2021.09.26.461864

**Authors:** Rachid Ramadan, Hartmut Geyer, John Jeka, Gregor Schöner, Hendrik Reimann

**Affiliations:** Institute for Neural Computation, Ruhr University Bochum, Bochum, Germany; Department of Kinesiology and Applied Physiology, University of Delaware, Newark, USA; Robotics Institute, Carnegie Mellon University, Pittsburgh, PA, USA

## Abstract

Existing models of human walking use low-level reflexes or neural oscillators to generate movement. While appropriate to generate the stable, rhythmic movement patterns of steady-state walking, these models lack the ability to change their movement patterns or spontaneously generate new movements in the specific, goal-directed way characteristic of voluntary movements. Here we present a neuromuscular model of human locomotion that bridges this gap and combines the ability to execute goal directed movements with the generation of stable, rhythmic movement patterns that are required for robust locomotion. The model represents goals for voluntary movements of the swing leg on the task level of swing leg joint kinematics. Smooth movements plans towards the goal configuration are generated on the task level and transformed into descending motor commands that execute the planned movements, using internal models. The movement goals and plans are updated in real time based on sensory feedback and task constraints. On the spinal level, the descending commands during the swing phase are integrated with a generic stretch reflex for each muscle. Stance leg control solely relies on dedicated spinal reflex pathways. Spinal reflexes stimulate Hill-type muscles that actuate a biomechanical model with eight internal joints and six free-body degrees of freedom. The model is able to generate voluntary, goal-directed reaching movements with the swing leg and combine multiple movements in a rhythmic sequence. During walking, the swing leg is moved in a goal-directed manner to a target that is updated in real-time based on sensory feedback to maintain upright balance, while the stance leg is stabilized by low-level reflexes and a behavioral organization switching between swing and stance control for each leg. With this combination of reflex-based stance leg and voluntary, goal-directed control of the swing leg, the model controller generates rhythmic, stable walking patterns in which the swing leg movement can be flexibly updated in real-time to step over or around obstacles.

## 1 Introduction

Walking is one of the most common movements humans perform every day. Walking consists of putting one foot in front of the other while moving the body forward. Most of the time walking does not require attention. But when walking in complex terrain, we are able to precisely step to suitable locations. When someone bumps into us, we are able to modify our normal movement pattern to maintain upright balance. In these situations, we are able to quickly and smoothly transition to conscious control of the usually largely habitual walking movement. The motor control of walking as a movement that is usually habitual and not consciously controlled, sometimes voluntary and goal-directed, and often somewhere in-between is currently not well understood. In this paper, we present a neuromechanical model for generating walking movements that is capable of covering the whole range of walking movements between these two poles.

### 1.1 Human Walking as a Voluntary Movement

Human movement shows amazing flexibility. We can perform a wide variety of tasks that require different movement patterns and coordination between body parts. Meaningful tasks usually require us to move a body part or tool to a goal position, such as the finger to a button or a screwdriver to a screw. Many tasks also contain additional requirements for timing or force, e.g. catching a ball in the air or hitting a nail with a hammer. The human nervous system routinely solves complex movements tasks in situations that it never specifically encountered before, using sensory information to generate a movement plan and update it during execution.

Humans are able to flexibly modify the basic pattern of their gait cycle during walking [106, 1]. At a high level, a walking movement pattern can be quantified by variables like speed and heading direction, and the length, width, duration and frequency of steps, typically referred to as gait parameters [66]. Humans can generally choose these parameters as desired. They can change direction, walk fast or slow, with narrow or wide steps and a slow or fast pace, etc. [45]. In addition to this high-level flexibility of gait patterns, humans are also able to choose how exactly they perform each low-level limb movement. Stepping to a fixed location in a fixed time can be performed with a variety of trajectories for the swing foot. We can swing the foot higher to step over an obstacle, or closer to the stance leg to step around an object. We can also choose to walk with bent knees, with the foot rotated in or out, or tip-toe by limiting ground contact to the balls of the foot and keep the heels up.

### 1.2 Stability and Upright Balance

One aspect of moving a body part to a target is the ability to confine movement to only the desired body part, while keeping the rest of the body stable and un-moving. Pushing a button requires not only muscles along the arm and shoulder to move the finger to the button, but also muscles along the trunk and legs to stabilize the rest of the body, so that the contact force at the finger results in moving the button in rather than the body away [129]. Muscle activation measurements reveal that when initiating such a manipulation movement while standing, muscles along the legs and trunk that stabilize the body activate earlier than muscles along the shoulder and arms that move the arm [5]. Stability is an integral part of the motor system that is integrated into the movement plan at all stages [10].

Stability is especially important for the upright body as a whole. When the body is upright during standing or walking, failure to stabilize it properly can lead to a fall, resulting in impact with the ground and serious injury. Yet for walking, “not moving” is not an option. We cannot keep parts of the body static relative to the environment, because locomotion of the whole body to a different place is the functional goal. The task for the nervous system is to generate a stable movement pattern for the whole body, transporting it with a relatively constant velocity from one point to another, while keeping movements in other directions to a minimum. To solve the main task of locomotion, the legs need to generate forces against the ground, initially to accelerate the body in the direction of travel and reach a steady state of motion, then to regulate the body movement around the steady state movement pattern and correct deviations from it. To prevent falls, the legs need to generate vertical forces that keep the body mass at a certain height, and also horizontal forces that regulate the body movement in the direction orthogonal to the direction of travel. Both of these requirements need to be combined into a cyclical pattern of moving one leg ahead in a step while supporting the body weight with the other one, then shifting weight and the role of the legs.

### 1.3 Habitual Control

Despite the flexibility to choose from a large range of walking patterns and movements, normal human walking is usually highly repetitive, with few variations. Humans will generally choose a walking pattern and then adhere to it for longer stretches of time, with gait parameters relatively stable on a time scale of minutes [22]. One factor driving this long-term stability of walking patterns is energy efficiency. The “cost of transport” of using metabolic energy to move from one place to another depends on the walking speed, with large cost at high and low speeds, and lower cost at medium speeds [90]. Humans usually choose to walk near the speed where this metabolic cost of transport is minimal [90, 11, 109]. A second factor affecting the choice of gait pattern is balance [8, 91]. Walking with increased step width increases the base of support during double stance, so the body is passively more stable [26]. But higher step width also leads to larger average displacement between the body center of mass and the stance foot during single stance, increasing the lever arm of the gravitational force pulling the body down, and thus the muscle forces required to counter gravity and keep us upright. Higher muscle forces require more metabolic energy, so there is a trade-off between balance and metabolic cost, where gait patterns that are more stable are also less efficient [25]. Balance is also actively maintained by changing the foot placement relative to the average gait pattern based on the current state of the body in space [123, 12, 93]. This active control of foot placement aggregates high-level sensory information about the body in space from the visual and vestibular and proprioceptive systems [84] and maps it to changes in foot placement. This mode of control is neither reflex-based in the narrow sense nor voluntary or conscious, but similar to online updating to a new target during a reaching movement [101].

The choice of gait pattern is different across different groups of people. Older people tend to walk more slowly [82, 94, 86]. People with Parkinson’s Disease tend to take short, shuffling steps [48]. People with Cerebral Palsy often swing their legs out to the side much more than typical [110]. While there are reasonable explanations for some of these gait pattern changes, the underlying causes are often not well understood. One reason for this limited understanding is the complexity of the problem. Walking is a biomechanically complex motor pattern with many moving parts [79]. The concrete choice of motor pattern depends on many different factors, including metabolic energy cost, avoiding muscle fatigue, stability and control of upright balance, and external constraints such as obstacles and the condition of the walking surface [56, 87, 121, 44, 72, 109, 58]. While motor control of walking is largely sub-conscious, cognitive processes also play a role, and secondary tasks during walking have routinely been shown to affect gait parameters and balance control [72].

### 1.4 Modeling Walking Control

To understand the interactions between different factors that drive the choice of walking movement pattern, we need a computational model that includes all factors of interest [3, 21]. Such a model allows us to manipulate individual factors and observe the resulting changes in the walking pattern directly in simulation studies [94]. Existing neuromechanical models of walking largely focus on the generation of rhythmic movement patterns and balance control. The rhythmic movement patterns are either generated by neural oscillators [113, 120] or by a finite state machine switching between different movement states depending on ground contact [35, 31]. These existing models have some degree of flexibility. Some models can walk at different speeds [114, 105, 120], change direction [120], and step over obstacles [112, 105]. This can be achieved by re-parameterizing a model, essentially optimizing a large number of neuromechanical parameters to walk at a range of different speeds, and then switching between these parameter sets, or interpolating between them, to change speed during walking [105, 120, 23]. Another approach is to modulate the central neural drive of a model to oscillate faster [114, 120]. Similar techniques can be used to step over obstacles, either increasing the gain between the central oscillator and the flexor muscles of the swing leg hip and knee [112], or the target flexion angle for a reflex at the same joints, with similar effect [105]. These approaches generally provide solutions for one specific problem, e.g. walking at different speeds or stepping over an obstacle, but do not generalize directly to related problems, such as walking at different cadences or stepping around an obstacle, rather than over it. Humans, in contrast, are not only capable of flexibly modulating gait parameters or the path of the swing foot, but can spontaneously walk in novel patterns, which they never used or observed before.

Our goal is to develop a neuromechanical model of walking that shows a similar degree of flexibility as humans, in that it can generate any desired walking pattern. We postulate that the key limitation of current walking models is that they are almost completely spinal, and lack cortical motor planning and control. These high-level features are usually studied as part of upper extremity reaching movements [49, 98, 20]. Some researchers have pointed out the duality of steps as (i) part of a cyclical movement pattern of the whole body for locomotion and (ii) a reaching movement with the foot [97, 96, 104, 75, 7]. Experimental evidence indicates that stepping movements during walking are generated rhythmically using low-level, reflexive structures [77, 47, 133, 107]. On the other hand, these movements can be precisely and efficiently modulated by high-level influences when desired, e.g. to step to a specific target or around an obstacle [135].

Here we present a model extension that attempts to bridge this gap between existing neuromechanical models of walking and the ability to plan and execute voluntary movements with the leg. The key innovation in our model is an explicit movement plan for the swing leg on task level. The high-level movement plan is executed by transforming the planned movements into descending commands that integrate with the low-level, reflex-based control architecture of the spinal cord, using internal models to account for dynamic interaction forces and properties of the muscles and spinal reflexes. For the stance leg, we use an existing solution of dedicated spinal reflex modules that generate the appropriate muscle activation with minimal high-level input [105]. We show that this model is able to generate voluntary swing leg movements, and to integrate these swing leg movements into a rhythmic walking pattern, modulated by high-level feedback to maintain upright balance.

## 2 Methods

The model spans multiple levels, across high-level movement planning and coordination, spinal reflex arcs, muscle physiology and skeletal biomechanics. Figure 1 provides an overview. A finite state machine organizes the model and switches between swing and stance phase control for each leg. In the supraspinal layer, a volition module represents task-level movement goals, a planning module generates motor plans to reach the goal state and a balance module updates the movement plan based on real-time sensory feedback about the body in space. An internal model then transforms the high-level motor plan into descending motor commands that interface with the spinal cord to execute the planned movement. In the spinal cord, the swing leg is controlled by a generic stretch reflex that is modulated by the descending commands, while the stance leg control is purely reflex-based. Reflexes stimulate Hill-type muscles that actuate a biomechanical model.

**Figure 1:**
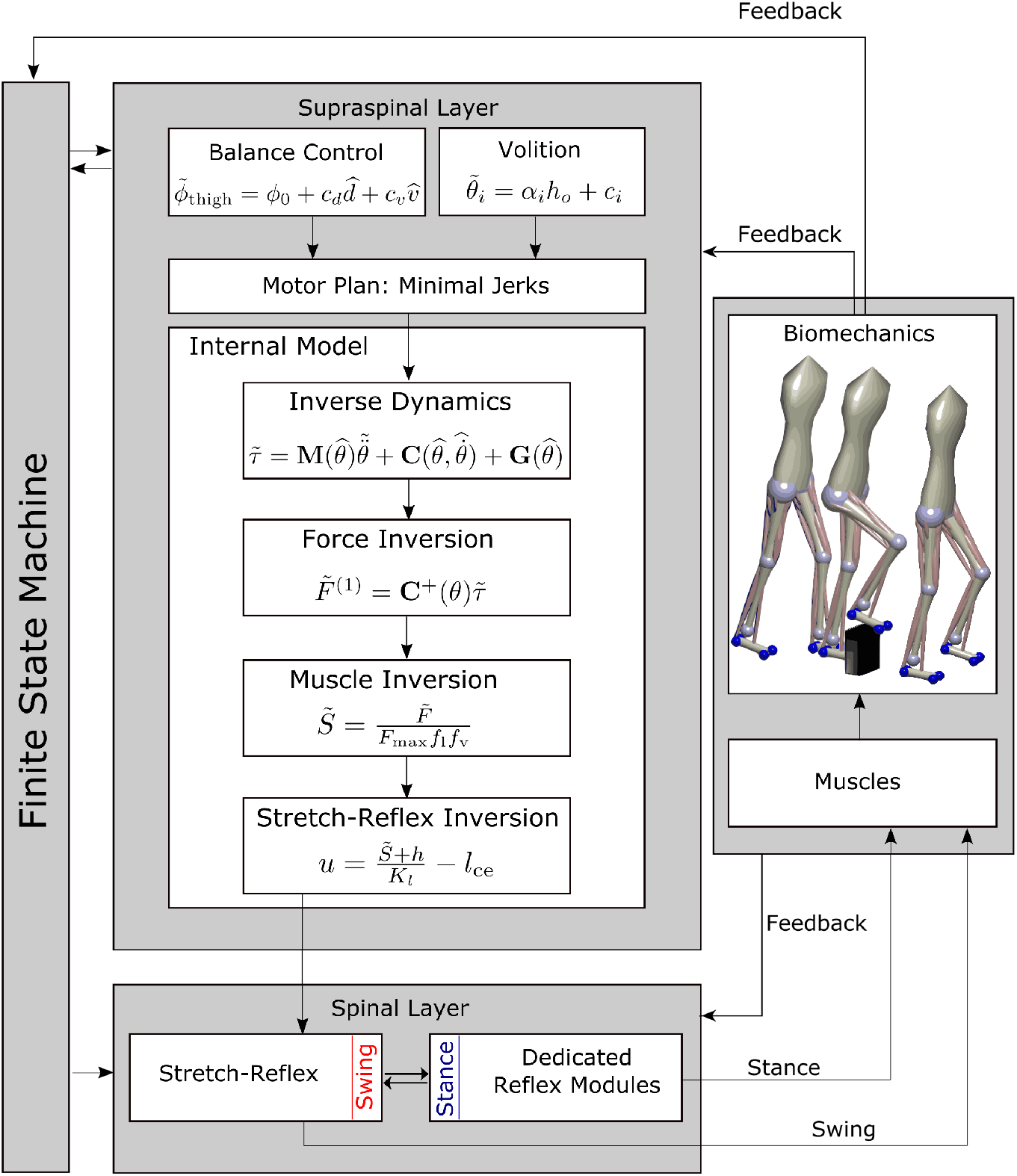
Overview of the model architecture. In the supraspinal layer, a balance control equation defines target joint angles for the swing leg at mid-swing and heel-strike. The target joint angles can be modified to perform volitional, goal-directed movements. A movement plan towards these target joint configurations is generated by minimal jerk trajectories that can be updated during execution. An internal inverse model comprising biomechanics, muscle moment arms, muscle activation properties and the spinal stretch reflex produces descending commands that realize the planned movement. The descending commands are integrated with the stretch reflex in the spinal layer. Stance leg control is realized with five dedicated reflex modules [105]. Reflex outputs are the applied to the biomechanical model that provides feedback to the controller. A finite state machine organizes switches between early swing phase, late swing phase and stance phase.

The key innovation here is the integration of the volition module in the supraspinal layer that prescribes movement goals with the other components. While the volition module itself is relatively simple, the main challenge in the development of this model was to integrate the task-level movement goals with the low-level spinal reflex control modules so that the resulting system can combine stable, repetitive walking movements with voluntary, goal-directed movements that solve tasks represented in the volition module. The finite state machine, balance control, spinal reflexes, muscle model and biomechanics are all modeled with standard solutions from textbooks or the literature. Each module is described in detail below. All methods were performed in accordance with the relevant guidelines and regulations.

### 2.1 High-level Control

#### 2.1.1 Behavioral organization

Walking requires the sequential execution of different movements for each limb, organized in a cyclical pattern [74, 28]. We organize the model behavior in three phases per leg, (1) early swing, (2) late swing and (3) stance. A finite state machine generates transitions between these phases based on sensory information of ground contact and internal timing. The early swing phase is initiated by the detection of ground contact of the contralateral leg (3 → 1). Transition to the late swing phase occurs after a fixed time of 0.3 s (1→ 2). The late swing phase lasts until ground contact is detected, leading to a transition to the stance phase (2 → 3). The finite state machine is adopted from [132] and functionally equivalent to one with four global states. When one leg is in the stance phase, the other leg is either in the early, or the late swing phase since ground contact detection of the swing leg triggers transitions between swing- and stance phase. Note that this finite state machine, including the transition from early to late swing phase based on explicit time with the transition time chosen in an ad-hoc way, is not necessarily physiologically plausible. Other models use more stratified sub-phases [81, 23].

During swing, the leg is controlled in a goal-directed way based on a movement plan (see Section 2.1.4 below). During stance, the leg is controlled in a purely reflex-based way (see Section 2.2.2 below).

#### 2.1.2 Volition

A goal for a voluntary movement of the swing limb is a desired configuration of the limb kinematics, represented by a vector of desired joint angles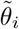.The tilde in 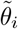 indicates that this is a desired state of the joint angle, in contrast to the actual joint angle *θ*_*i*_. We will use this convention of the tilde to denote desired states throughout the rest of the text. In principle, this goal configuration can be anything, and we will probe the generation of movements to randomly chosen configurations (see Section 3.1). For walking, the goal configuration for each movement phase must be appropriate to generate a stable gait pattern, and we use evolutionary optimization to find suitable configurations. During individual steps, the goal configurations can be modified to address specific tasks, such as obstacle avoidance (see Section 3.2 or balance control (see Section 2.1.3).

#### 2.1.3 Balance Control

Maintaining balance requires the integration of state feedback about the body in space into the movement plan. We use position and velocity feedback of the trunk center to update the desired target orientation of the thigh in space. Following [131], we use the control law

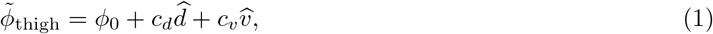

where *ϕ*_thigh_ is the desired orientation of the swing leg thigh, *ϕ*_0_ is a constant offset, 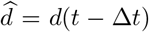 are the time-delayed horizontal displacement from the center of pressure (CoP) to the trunk segment center, 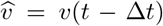 the time-delayed rate of change of that displacement and *c*_*d*_ and *c*_*v*_ are feedback gains. Equation 1 is applied independently for the sagittal and frontal plane orientation of the thigh. We then calculate target joint angles for each DoFs of the hip joint

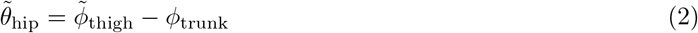

by subtracting the trunk orientation *θ*_trunk,world_ from desired thigh orientation, again separately in the frontal and sagittal planes. Note that if the target joint angle for the knee in the late swing phase is close to zero, the thigh angle will correspond closely to the swing leg angle in space, which is relevant for balance.

#### 2.1.4 Movement Planning

The swing leg is controlled in a goal-directed way according to a task-level motor plan. The task-level goal is a kinematic configuration of the swing leg, defined by the swing leg joint angles, combined with a target time at which the goal configuration should be reached. Goal configurations and target times are different for early and late swing phase and can be updated to maintain whole-body balance (see Section 2.1.3 above) or to generate specific voluntary movements. The leg will typically be far away from the goal configuration at the onset of each movement phase, and there is an infinite number of possible movement trajectories that will fulfill the task constraints. Human movements are generally smooth and avoid unnecessary spikes in force and acceleration, and a standard way to plan such movements are minimum jerk trajectories [43].

For a given combination of initial state

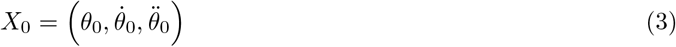

and goal state

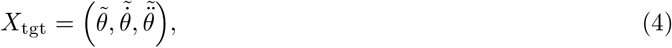

of joint angles, velocities and accelerations for a single joint angle *θ*, and a movement duration *T*, the minimum jerk trajectory is a 5th-order polynomial

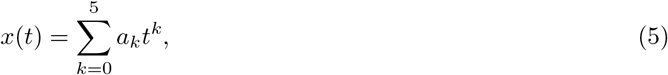

with parameters *a*_*k*_ that fulfill the constraints

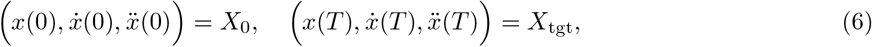

which can be computed analytically depending on *T, X*_0_ and *X*_tgt_. We use a version of the minimal jerk approach that allows changes in target states and time before the movement is complete. For every moment in time *t*, we regard the current state estimate

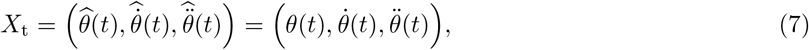

as the initial state of a new movement and compute parameters *a*_*k*_(*t*) such that in the remaining time (*T* −*t*), the movement reaches the target state *X*_tgt_. From the resulting parameters *a*_*k*_(*t*), we compute the jerks

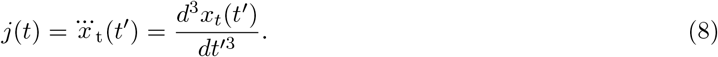

For very small remaining movement times *T* − *t <* 0.03 s, we stop updating the motor plan. Integrating these jerks over time yields a desired joint acceleration

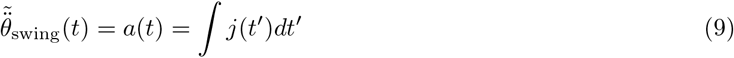

to be realized with descending motor commands.

The tilde indicates that this is a planned, or desired, joint acceleration, in contrast to the actually realized joint acceleration that will be a combination of active and passive muscle-tendon forces, gravity, ground reaction forces and interaction torques. We will keep this convention to use a tilde to indicate planned or desired values for a variable from here on. By applying this procedure of updating the planned trajectory based on the estimated state during the entire movement, we are able to adapt the initial minimal jerk trajectory to account for any external or internal perturbation and correct the resulting errors.

We use this procedure to generate a minimum jerk trajectory for each degree of freedom in the swing leg that moves the leg to the target configuration in the given time. The target joint angles for early swing and late swing are part of the parameters set that is determined by evolutionary optimization (see Section 2.4 below).

#### 2.1.5 Transformation into Descending Motor Commands

The motor plan is represented by a minimum-jerk trajectory that moves the joint configuration to the desired state in the remaining time (see Section 2.1.4 above). At each point in time, this planned trajectory defines a vector of desired joint accelerations 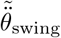 for the swing leg. Executing the motor plan means realizing these planned joint accelerations. Here we describe how this vector of desired joint accelerations is transformed into a descending motor command that executes the motor plan. We solve this problem using inverse models of the biomechanics, muscle force dynamics and spinal reflex arcs, with simplifying assumptions.

##### Inverse Dynamics

The biomechanical Equation of Motion (17) relates joint accelerations to joint torques. We augment the planned vector of joint accelerations for the four degrees of freedom in the swing leg by zeros [103] in the components for the stance leg and the six free-body degrees of freedom of the trunk to get a vector of joint acceleration for the full 14-DoF model

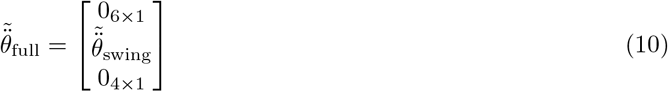

and use an Equation (17) to get a planned joint torque vector

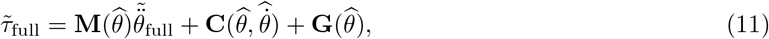

where 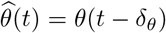 and 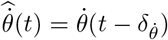 are time-delayed sensor estimates of the body configuration and rate of change. We then take the swing leg components 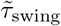 of

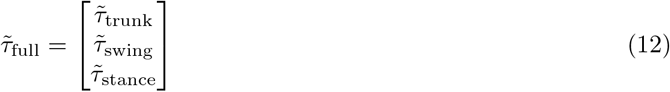

as the desired joint torques for the swing leg that will execute the motor plan. We implement Equation 11 using the Inverse Dynamics block in Simulink.

##### Muscle Moment Arm Inversion

In order to obtain a set of muscle forces 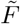 that generate the desired joint torques 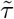, we use the Moore-Penrose pseudo-inverse [76] of the moment arm matrix *C* to get

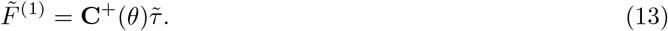

The resulting force vector 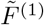, however, can contain negative forces, which cannot physically be generated by muscles. Instead of using a computationally intensive solution like the non-negative least squares [64], we use an iterative approximation. We separate the negative part of the resulting forces 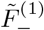, consisting of the muscle forces with negative signs from the positive part of the forces 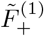 and compute the joint torques produced only by the negative forces 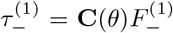. We then apply Equation 13 again on these torques, getting 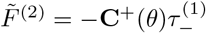, which will also contain both positive and negative forces. Iterating this procedure leads to progressively smaller remaining negative forces 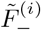.

We apply this procedure for a total number of 7 iterations and sum up all positive forces to obtain 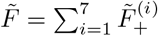 as a force vector that generates the joint torques 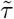 with an root-mean-squared error of below 3 *N* per joint for a usual stepping movement.

##### Inverse Muscle Model

The force generated by a muscle depends on its activation level and its current length and velocity. We compute the activation needed to generate the desired muscle force 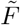 by inverting the muscle model, with some simplifications. We neglect the low pass filtering of the muscle activation which models the excitation-contraction coupling, setting 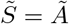. We approximate the total muscle force *F*_*se*_ with the force of the contractile element *F*_*se*_, neglecting the contributions of the passive buffer and parallel elements. This is reasonable because the buffer and the parallel element are active only when muscles are extensively stretched or compressed, which is usually not the case during walking.

We then invert Equation 19 to calculate the neural stimulation 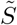 needed to generate the desired muscle force as

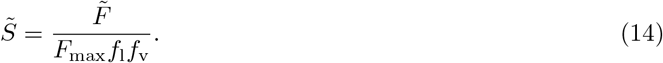

All terms here are 22-dimensional vectors, with one component per muscle, and the operations are executed element-wise.

##### Spinal Stretch Reflex Modulation

The descending commands from the high-level motor areas have to interface with the reflex arcs in the spinal cord to generate muscle activation levels that will execute the planned movement. Described in detail in Section 2.2 below, we assume that the descending command both *(i)* directly creates muscle activation leading to contraction and *(ii)* shifts the reference point of the spinal stretch reflex to a new location corresponding to the contracted state. We solve Equation 16, which models this behavior, to calculate a descending motor command

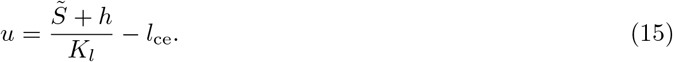

Note that while we neglected the velocity term in the stretch reflex 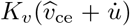 here, it is this velocity-dependent term that will initially create the direct muscle activation, determined by 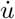. This descending motor command *u* will interact with the spinal stretch reflex to generate the desired muscle activation 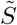 that executes the motor plan.

### 2.2 Spinal Control

Spinal control consists of neural feedback loops, i.e. feedback laws that generate neural activation proportional to low-level proprioceptive signals about muscle length, velocity or force, modulated by descending commands on a slower time-scale. We treat control of the leg during swing separately from the control of the leg during stance. While the swing leg is controlled by a combination of descending commands and a generic stretch reflex, the stance leg is controlled by specialized reflex modules that implement a specific function.

#### 2.2.1 Swing Leg

During swing, the neural stimulation for each muscle is generated by a generic stretch reflex

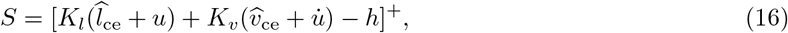

where 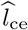 and 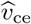 are proprioceptive signals from muscle spindles that estimate the stretch and stretch rate of change of the muscle contractile element, *K*_*l*_ and *K*_*v*_ are gain factors, *h* is the resting level activation of the *α*-motorneuron and *u* and 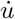 are the descending motor command and its rate of change.

Note that the descending command *u* acts as a threshold for the reflex loop and the rate of change 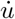 is used for relative damping. When the descending command *u* increases to contract the muscle, both *u* and 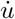 will increase initially, generating a stimulation burst that is mostly driven by the rate of change 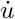. While formulated as a single stretch reflex with relative damping here, this is functionally equivalent to a formulation where the *α*-motorneuron activation level is determined by a sum of a spinal stretch reflex and a descending motor command, as used in other models [27, 33, 35, 57, 13].

This principle of modulating a generic stretch reflex with a descending motor command leads to the flexibility to execute motor plans for goal-directed movements via appropriately chosen descending commands, combined with the robustness of a stretch reflex that provides a level of postural stability to the muscle-joint system in situations where it is not part of a goal-directed movement.

#### 2.2.2 Stance Leg

During stance, the leg is controlled by purely spinal mechanisms, without modulation by descending motor commands and without the flexibility to execute goal-directed movements. Proprioceptive information from different muscles and joints is mapped to proportional muscle activation in a set of dedicated neural control laws that implement specific functions, organized in five modules following [105].

Functionally, the modules (1) generate compliant, spring-like leg behavior, (2) prevent knee overextension, (3) keep the trunk upright, (4) compensate interaction torques from swing leg movements and (5) dorsiflex the ankle joint to prevent hyperextension.

Proprioceptive feedback mechanisms incorporated in the model include positive force feedback provided by the Golgi tendon organs, length feedback from the muscle spindles, and feedback about muscle activations and joint states. The stretch reflex, as modeled in 16, is not used during the stance phase. Please refer to [105] for details.

### 2.3 Muscoskeletal Mechanics

#### 2.3.1 Body Model

The body model represents a person of 180 cm height and 80 kg weight. It is composed of seven body segments, eight degrees of freedom (DoF) and 22 muscle-tendon units (MTU). Body segments comprise two thighs, shanks and feet, and a trunk segment that represents the entire upper body, including head and arms [105]. Revolute joints link the body segments with two DoFs at each hip (pitch and roll), one DoF at the knees (pitch) and one DoF at the ankles (pitch). The equation of motion

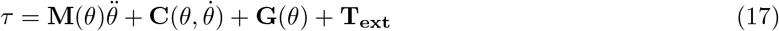

relates joint torques *τ*, gravitational torques **G** and external torques **T**_**ext**_ to joint accelerations 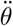, where **M** represents the mass matrix and **C** the velocity dependent terms. Joint accelerations 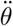 and torques *τ* are 14-dimensional vectors, with the eight internal DoFs and six free-body DoFs for translation and orientation of the trunk segment. Note that the six free-body DoFs of the trunk are un-actuated. Geometry and inertia of the body segments are adopted from [105].

Each leg is actuated by eleven Hill-type MTUs that are either mono- or biarticulary (see Section 2.3.2 below for details). Nine MTUs actuate the three pitch joints (hip, knee, ankle) and two MTUs actuate the hip roll joint. Pitch joint muscles model the lumped hip flexors, glutei, hamstrings, rectus femoris, vasti, biceps femoris short head, gastrocnemius, soleus and tibialis anterior. Roll joint muscles represent the lumped hip adductors and hip abductors. Muscles forces translate into joint torques via state-dependent moment arms that are adopted from [105], via

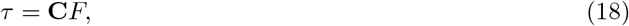

where *F* is the 22-dimensional vector of muscle forces and *C* is the 14 *×* 22 matrix of moment arms.

#### 2.3.2 Muscle-Tendon Units

Each muscle tendon unit (MTU) is composed of a parallel element (PE), a buffer element (BE), a contractile element (CE) and a serial elastic element (SE). We provide an overview here and refer the reader to [31] for details. The contractile element is the actual active muscle element. It is innervated by the *α*-motorneurons and exerts the force

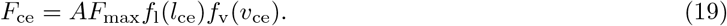

Here, *F*_max_ is the maximum isometric force, *f*_l_(*l*_ce_) and *f*_v_(*v*_ce_) are the force-length and force-velocity relationships and *A* is the muscle activation level. The serial element models the tendon and applies the generated forces *F*_*se*_ to the body. The parallel element passively prevents the muscle from being stretched extensively and exerts a force *F*_pe_(*l*_ce_) after the muscle lengths exceeds a certain maximal length. In contrast, The buffer element is a passive element that prevents the muscle from being compressed too much. It generates the force *F*_be_(*l*_ce_) only after the muscle length shortens below a certain minimal length. Muscle activation *A* is modeled as a first-order low-pass filtered copy of the neural stimulation *S* representing the *α*-motorneuron output

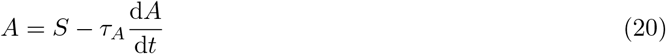

where *τ*_*A*_ is a time constant [33]. The total force *F*_mtu_ generated by a MTU is given by

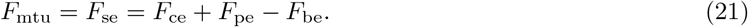

#### 2.3.3 Ground Contact Forces

Ground contacts at each foot are modeled with four contact points, two at the heel and two at the front of the foot, with a lateral displacement of 5 cm between the two points at the heel and 10 cm at the front. We compute ground reaction forces by using the inbuilt MATLAB Spatial Contact Force block. Contact parameters are chosen to simulate an asphalt surface.

### 2.4 Parameters and Tuning

The model contains a large number of parameters for different components of the model. Some of these parameters are constrained by the neurophysiological literature and set to constant values based on estimates. To determine the other parameters, we use an evolutionary optimization algorithm similar to the one used in[105], based on the covariance-matrix adaptation technique [36], using the cost function

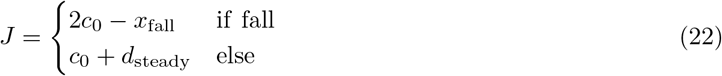

The first part of the cost function generates basic walking without falling and the second part generates steady locomotion. The constant *c*_0_ = 10^3^ is a normalization factor and *d*_steady_ measures the “steadyness” of the gait. We calculate *d*_steady_ as

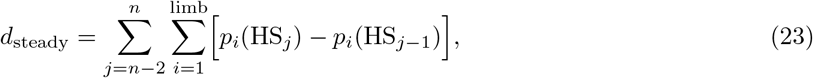

with *p*_*i*_ being the relative Cartesian position of the *i*-th limb and HS_*j*_ being the *j*th-last left heelstrike.

We optimize a total amount of 52 parameters. The same set of parameters is used for all experiments described in the results section.

### 2.5 Simulation Studies

#### 2.5.1 Swing Leg Movement

We evaluate the ability of the model to plan and execute voluntary movements with the swing leg a series of simulation studies. In each of the studies, we demonstrate that the control of voluntary movement works and the limb follows the movement trajectory as planned. To isolate the swing leg and remove balance control as a factor for these stimulation studies, we passively stabilize the trunk segment by fixing its position in space. In each of the experiments, the model performs reaching movements with the foot to different target locations that are defined either in joint space or as positions for the ankle and transformed into joint space using the inverse kinematics solution in the MATLAB RigidBodyTree toolbox. The movement plan in joint space from the current to the target configuration is then generated as described in Section 2.1.4 above. We perform a total number of three experiments. In the first simulation study, we explore whether the system can generate voluntary, goal-directed movements in isolation. The foot performs a sequence of center-out reaching movements to different target locations that cover a large portion of the workspace. In the second simulation study, we test whether the system can generate stable oscillatory patterns. The four joints simultaneously follow sinusoidal trajectories that with a frequency of 1 Hz. Amplitudes of the sinusoidal trajectories were chosen to roughly capture the ranges of a stepping movement. In the third simulation study, we explore whether the system can successfully reach to different points in the work space. The leg performs a total number of 1100 randomized movements. Each of the movements is drawn from a uniform distribution over the interval from 15–85% of the joint range of motion for each joint. Joint configurations that require muscle lengths smaller than the slack length of the corresponding muscle are excluded. For 11 different movement times ranging from 100 ms to 1000 ms, we simulate 100 randomized movements each.

#### 2.5.2 Obstacle Avoidance

We use an obstacle avoidance task to assess the ability of the model to integrate flexible swing leg movements control of upright balance during walking. We test two different avoidance strategies, (1) lifting the swing leg to step over an obstacle and (2) shifting the swing leg sideways to step around an obstacle. To avoid an obstacle, we adjust the movement plan for the early swing phase by updating the target joint angles 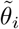 for the early swing phase based on the obstacle position and size. We used a linear mapping

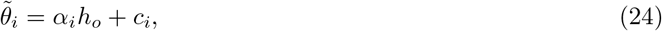

to determine the target joint angles, where *h*_*o*_ is the obstacle extension, i.e. the height for sagittal and the width for medial-lateral avoidance, including a security margin. The joint index *i* ranges over the ankle, knee and hip flexion degrees of freedom for sagittal and the hip abduction joint for medial-lateral avoidance, and *c*_*i*_ is a constant offset. We determined these parameters in an ad-hoc manner based on a few sample movements with hand-fitted values. To avoid the obstacle, the model changes the target joint angles for the early swing phase according to Equation 24, defining an alternative trajectory that achieves obstacle avoidance. After mid-swing, foot placement and balance recovery is controlled by the usual balance-control strategy described in Section 2.1.3.

#### 2.5.3 Direction and Speed Control

In order to evaluate the ability of the model to change movement direction and speed, we exploit the passive properties of the model by implementing ad hoc control laws. Change of walking direction is archived by temporarily adding a constant offset value to the hip roll target angle that causes a weak destabilization of the model and induces a rotational slipping of the stance foot. We add the constant offset to the hip roll target value when the horizontal orientation of the trunk segment lies outside a desired interval around the target orientation. We generate change in the movement speed by varying the target orientation of the trunk in the spinal reflex module for trunk stabilization (see Section 2.2.2 and [105]). A increase of the target orientation of the trunk leads to a larger trunk lean that results stronger gravitational accelerations.

## 3 Results

The model generates stable walking behavior with a movement speed of about 1.3 m/s. The walking pattern roughly matches human data. Figure 2 compares joint angle trajectories across one gait cycle averaged over a 100 s walk to human walking data from a public data set [28]. The human data is from N=24 healthy young participants (10 female, age 27.6 *±* 4.4 years, height 171.1 *±* 10.5 cm, and mass 68.4 *±* 12.2 kg) walking overground at their self-selected, comfortable speed. All reported root-mean-squared-errors are normalized to the range of motion. Panel A shows hip flexion angle for the model (blue) with human data (orange). The overall shape of the model trajectory matches human data. At about 90 % of the gait cycle, the model flexes the hip more strongly than the average experimental data. The model movements are also less smooth than the experimental data. The root-mean-squared-error of the hip flexion joint amounts 0.085.

**Figure 2:**
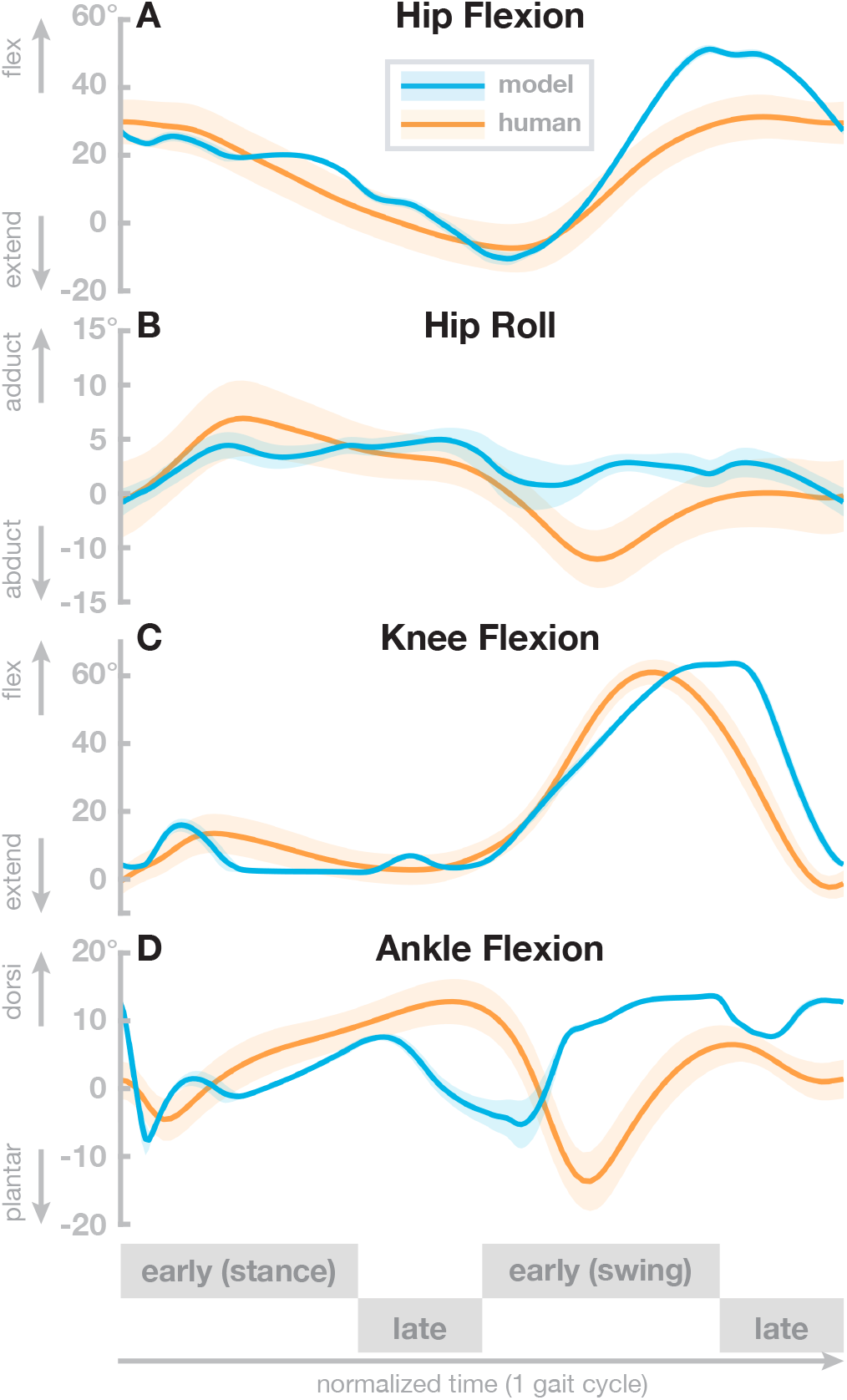
Comparison with human data. Averaged hip pitch, hip roll, knee and ankle joint angle trajectories for human data (orange) and model data (blue). The model data are averaged over 100 seconds of steady state walking. Human data are taken from [28]. Solid lines are means and shaded areas show standard deviation, across participants for the human data and across gait cycles for the model.

Panel B shows the hip adduction angle. Here, the overall shape of the model data differs from human data. The model trajectory is less smooth and has less overall range of motion throughout the gait cycle. Note, however, that the hip adduction in humans is quite variable, and despite the structural differences, the model data lies within the confidence interval of human data during a large part of the gait cycle. The root-mean-squared error for the hip abduction joint is 0.49.

Panel C shows the knee flexion angle. The model generally follows the human pattern. During the stance phase, the model exhibits two sharp peaks while human data shows one wider peak in contrast. During swing, humans extend their leg a little earlier then the model does. We found a root-mean-squared-error of 0.18. Panel D shows the ankle flexion angle. The overall pattern of the model trajectories differs significantly from the human data. The ankle dorsiflexion peak is slightly after mid-stance, much earlier than in the human data. In swing, the model shows consistently higher dorsiflexion than humans. The root-mean-squared-error of the ankle dorsiflexion amounts 0.34. The model trajectories of all joints exceeds two standard deviations of the average human trajectories [37]. However, a fit within two standard-deviations is not expected since the model is not calibrated with human data.

In order to investigate the robustness of the models walking behavior, we exposed it to external perturbations in the form of force pulses of increasing strengths applied at the center of the trunk segment in different directions. Perturbations started at foot contact, lasted for 0.2 s, and were directed forward, backward, medially or laterally. Force amplitude was ramped up until the model failed to maintain balance after the perturbation, starting at 50 N and increasing in steps of 50 N. After the model fell, we decreased the step size to 5 N from the previous value, until it fell again. The maximal force the model was able to withstand without falling was 340 N for lateral, 305 N for medial, 165 N for forward and 130 N for backward pushes. Perturbation studies with comparable pushes have been applied to humans [65] and were considered to be moderate pushes that could be recovered within 3 steps.

### 3.1 Swing Leg Movement

In the first simulation study, we show that the model can perform individual reaching movements with the foot. The model performs a sequence of center-out reaching movements with the foot to twelve different target locations, followed by a return movement to the center location. Target locations were defined as positions for the ankle and transformed into joint space. The movement plan was then generated as described in section 2.1.4. The specific target locations were arbitrarily chosen to cover a large portion of the workspace resulting in path lengths between roughly 0.20–0.55 m. Each single movement segment had a duration of 0.5 s. Figure 3 shows the resulting movement paths of the ankle position in workspace for this sequence of reaching movements. The ankle always reaches the target positions reasonably well, with a maximum deviation of 0.12 mm from the desired target position at the end of each movement. The largest deviation from the planned path is at the start of the first movement, to the top right target, which is due to the muscles being initialized without tension.

**Figure 3:**
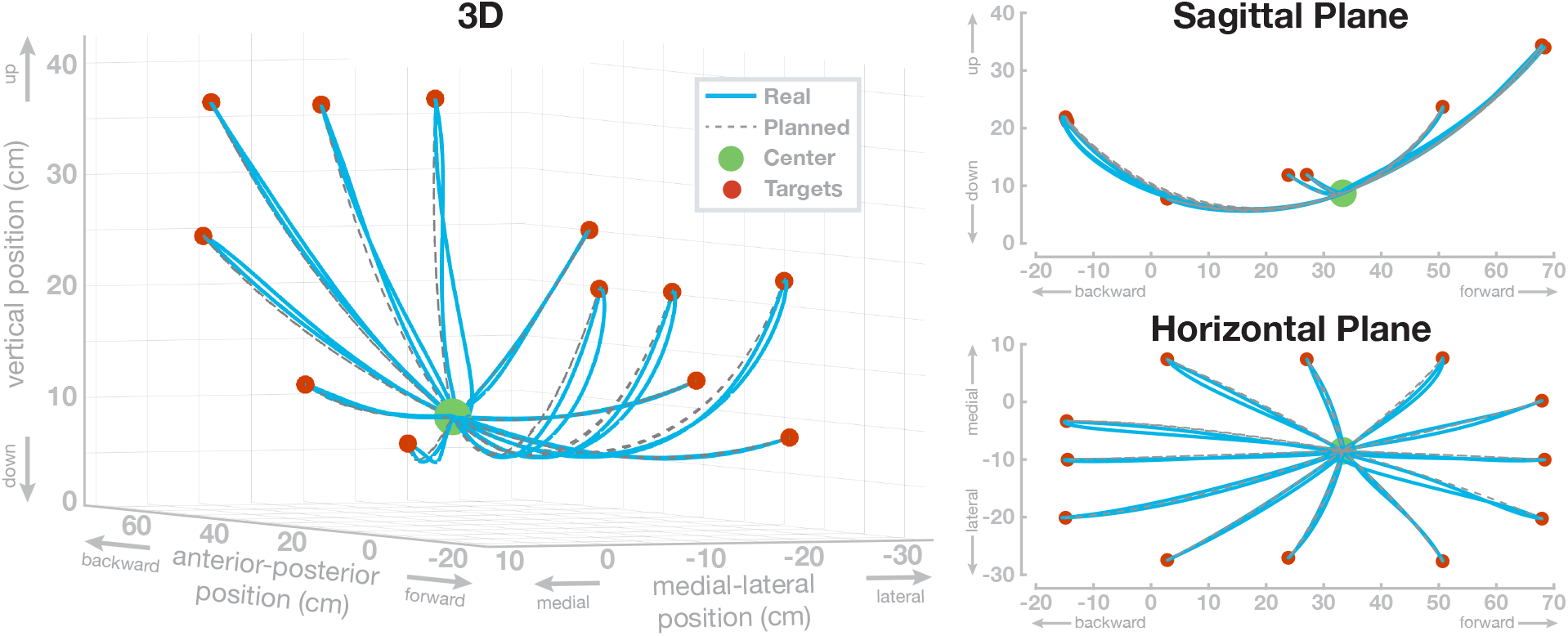
Swing leg ankle paths for a sequence of twelve center-out-return movements with passively stabilized trunk. The same movements are shown in three different views. (A) 3D perspective view, (B) side view of the sagittal plane and (C) top-down view of the horizontal plane. The dashed lines show the planned paths and the blue lines show the realized ankle paths.

The paths match the reference paths with an normalized root-mean-squared-error of 0.028 between the planned and the actual ankle position. To investigate a possible effect of movement extent on the error, we tested if the path lengths of the 24 individual movements are correlated with the non-normalized root-mean-squared errors of the Cartesian ankle position and found no significant correlation (*R* = 0.33, *p* = 0.11). Additionally, we tested if the path lengths are correlated with the non-normalized root-mean-squared-errors of the four individual joints. For hip flexion, we found a significant correlation (*R* = 0.71, *p <* 0.01). Correlations were not significant for hip abduction (*R* = −0.16, *p* = 0.45), knee flexion (*R* = 0.36, *p* = 0.08) and ankle flexion (*R* = 0.25, *p* = 0.18).

In the second simulation study, the model performs repetitive goal-directed movements between two points in joint space over 10 s. The four joints simultaneously follow sinusoid profiles with a frequency of 1 Hz. Figure 4 shows the resulting joint angle trajectories (solid lines) and the planned trajectories for each joint (dashed lines). The real joint angle trajectories fit the planned movement, with only the ankle joint showing more than minimal deviation of the real trajectories from the movement plan. The normalized root-mean-squared-error is 0.022 for the hip flexion joint, 0.018 for the hip abduction joint, 0.029 for the knee joint and 0.09 for the ankle joint.

**Figure 4:**
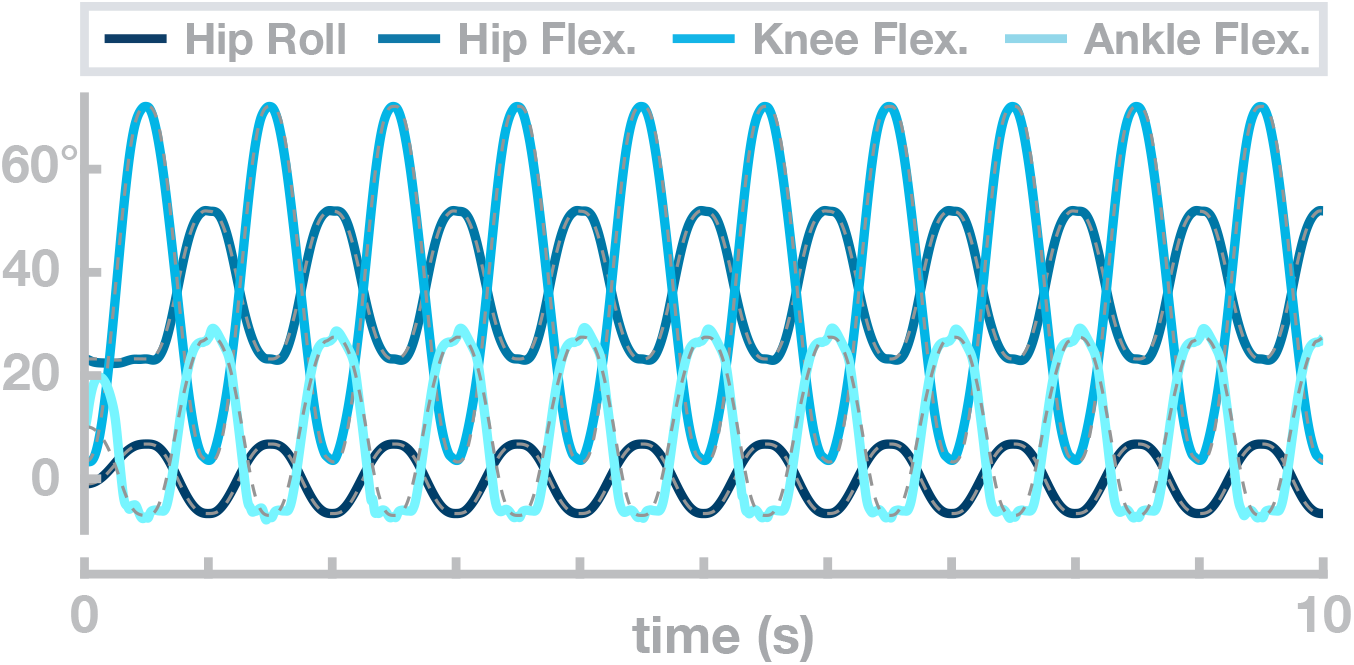
Example movement trajectories of the swing leg with a passively stabilized trunk. The dashed black lines show the planned movement trajectory and the blue lines show the realized joint angle trajectories.

In a third simulation study, we explore the flexibility of the model to generate goal-directed reaching movements with the foot between randomly chosen points in the joint space at a wide range of different speeds.

For each movement, the target configuration was drawn from a uniform distribution over the interval from 15–85% of the joint range of motion for each joint. Joint configurations that require muscle length below muscle slack lengths were excluded. We quantified performance as the normalized root-mean-squared-error between the actual and the planned trajectory. Figure 5 shows the average error for the different movement speeds in joint space. The error is high for movements faster than 0.3 s and reaches a minimum at 0.4 seconds. In the range that is relevant for walking [67], the error drops below 0.1 for hip and knee joints, and below 0.2 for the ankle joint. The root-mean-squared-error increases for slow movements because the minimum jerk planner corrects less strongly for errors early in the movement. We tested if the path lengths of the 1100 individual movements are correlated with the non-normalized root-mean-squared errors of the Cartesian ankle position and found no significant correlation (*R* = 0.05, *p* = 0.07). Furthermore, we tested if the path lengths are correlated with the non-normalized root-mean-squared-errors of the four individual joints and found no significant correlation for hip flexion (*R* = 0.05, *p* = 0.12), hip abduction (*R* = −0.01, *p* = 0.87) and knee flexion (*R* = 0.03, *p* = 0.29). For the ankle flexion the correlation was significant with (*R* = 0.06, *p* = 0.04).

**Figure 5:**
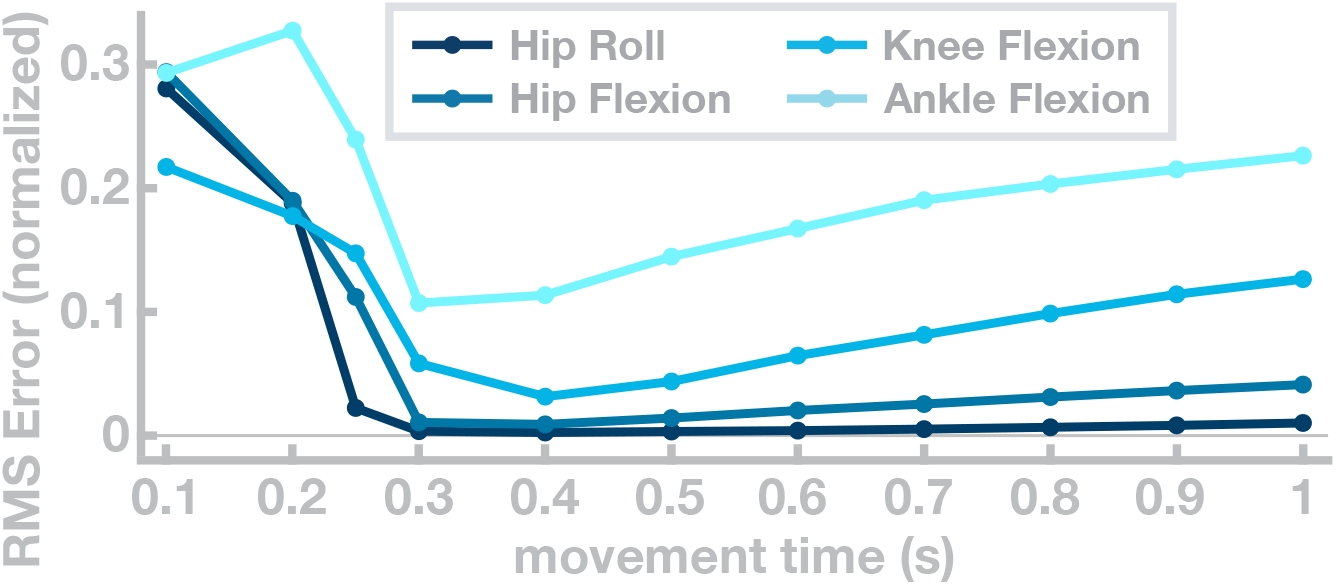
Joint-space error for single movements with different movement times. Each curve shows the root-mean-squared error the respective joint.

### 3.2 Obstacle Avoidance

We tested lifting the swing leg to step over an obstacle and shifting the swing leg sideways to step around an obstacle on obstacles of different sizes. An example of the model stepping over an obstacle is shown on the right in Figure 1. For sagittal avoidance, we simulated obstacles of 15cm, 20cm and 25 cm height. All three obstacles are successfully avoided and the model returns to the original gait within the subsequent step. Panel A in Figure 6 shows the difference between the balls of the foot relative to the normal movement with no obstacle, in the vertical direction, for these movements. The peaks of these difference plots show that in each movement the balls of the foot are successfully shifted upwards by the obstacle height, plus a safety margin. Note that these vertical positions are differences from the normal foot movement trajectory and the absolute vertical position of the foot is higher, around ≈ 10 cm at mid-swing.

**Figure 6:**
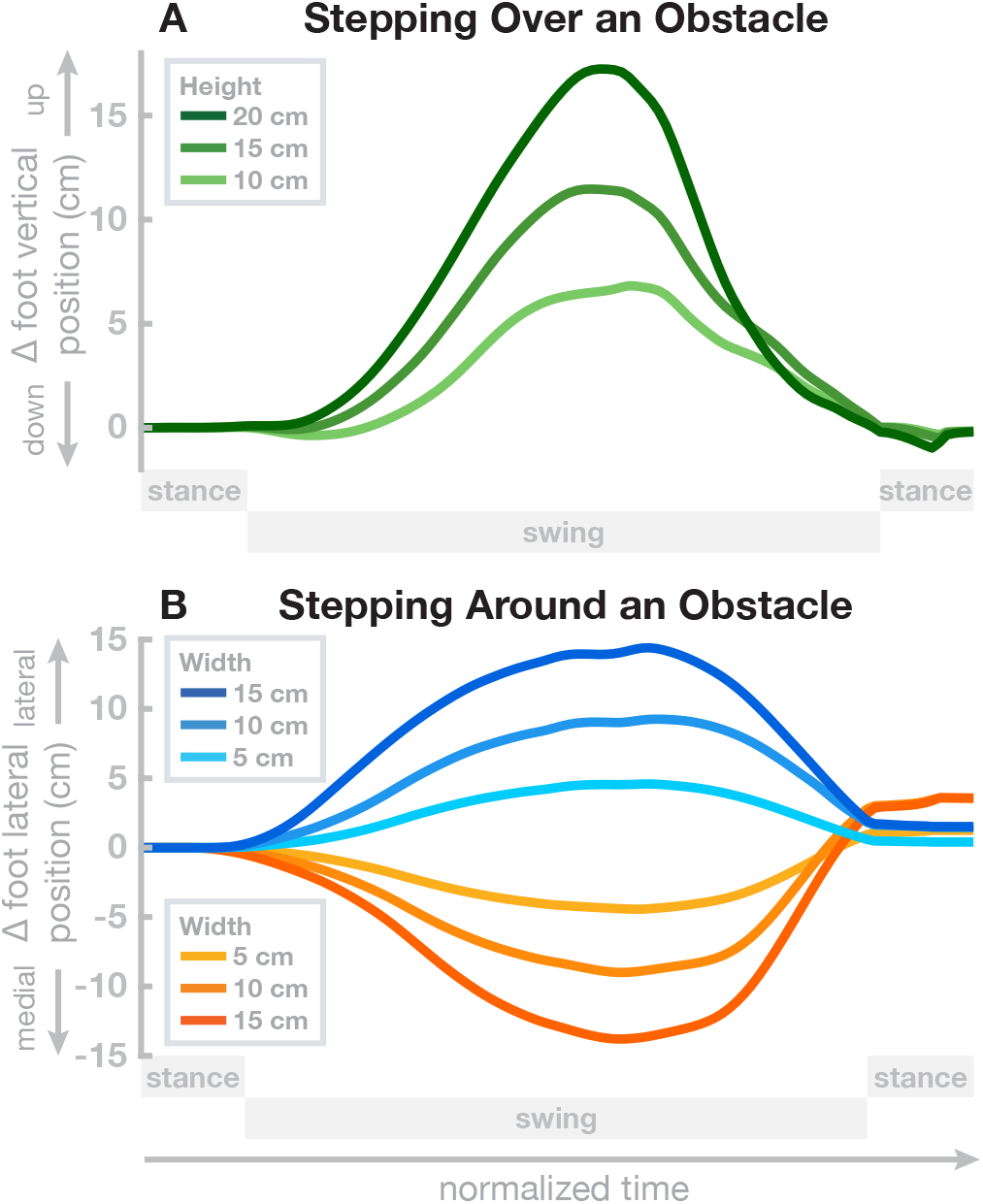
Avoiding obstacle during steady-state walking. Panel A shows the change from the normal trajectory without obstacle in the vertical direction when stepping over obstacles of varying height. Panel B shows the change from the normal trajectory without obstacle in the medial-lateral direction when stepping around obstacles of varying width, in either direction. Both panels show the movement from left heel-strike to push-off of the stance foot.

For medial-lateral avoidance, we simulated obstacles of 5 cm and 10 cm width, and both lateral and medial avoidance. Panel B in Figure 6 shows the differences between the balls of the foot relative to the normal movement with no obstacle, in the medial-lateral direction, for these movements. Similar to the sagittal avoidance, the peaks show that the movements are successfully shifted sideways by the desired amount corresponding to the width of the obstacle, plus a safety margin.

### 3.3 Direction and Speed Control

The model has a limited degree of flexibility to walk at different movement speeds and change direction.

Generally, increasing the trunk forward lean makes the model walk faster. To explore this relationship, we simulated 40 s of the model walking with 13 different random values for target orientation of the trunk, drawn from a uniform distribution between 6–8.5°. Figure 7 plots the resulting walking speed of the model in Panel A. Walking speed depends roughly linearly on the trunk, as shown by the linear fit (*R*^2^ = 0.9157). Panel B in Figure 7 shows how the stepping cadence varies depending on trunk lean for the same walking simulations. Interestingly, higher walking speeds are associated with lower cadences. This is opposite to what is observed in humans, where cadence tends to increase with walking speed in normal walking [80]. We interpret this as an indicator that the speed variations are not actively controlled, but rather emerge from the interaction of the trunk lean with balance control. Increased trunk lean leads to larger gravitational acceleration and higher speeds, which results in the balance control module increasing the target for the swing leg angle (Equation 1). This generates longer steps and decreases cadence.

**Figure 7:**
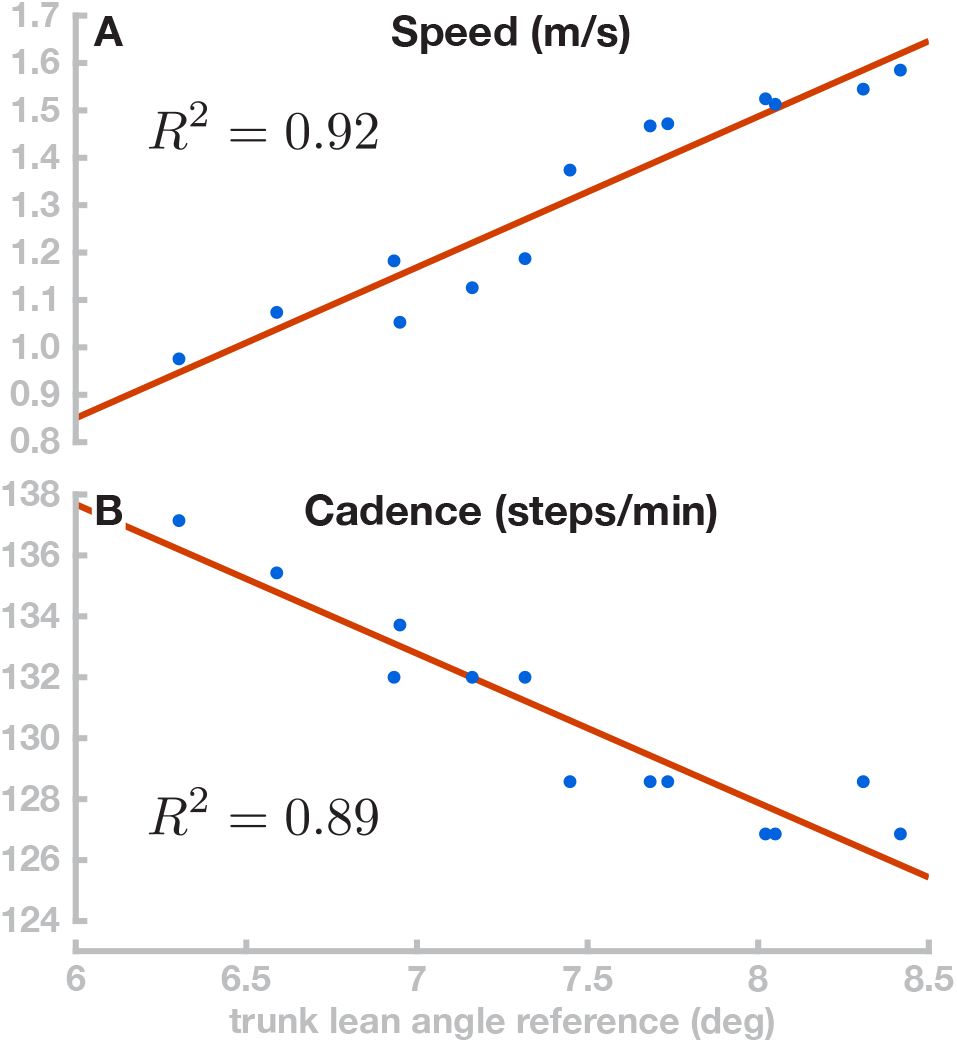
Velocity and cadence control. Panel A shows the relationship between the reference parameter for trunk lean and the resulting movement speed for 13 movements (blue dots) and the linear fit (red line). Panel B shows the effect of the trunk lean change on the walking cadence.

Although the model has no rotational degree of freedom at the hip, it is possible to change the direction of movement in a limited fashion. We use the ad hoc control law described in 2.5.3 and demonstrate the direction control scheme by simulating four walks with different target orientations of 0°, 15°, 30° and 45°, all starting at 0° and simulated for 100 s. Figure 8 shows the resulting walking patterns. For all four target orientations, the model approximately turns to the target orientation after about 20 m walking. However, this mode of direction control is not very stable and has clear limitations. The 15 deg and 30 deg movements turn away from the target orientation at about 16 meters of walking even though they reached the target orientation relatively fast after 12 m.

**Figure 8:**
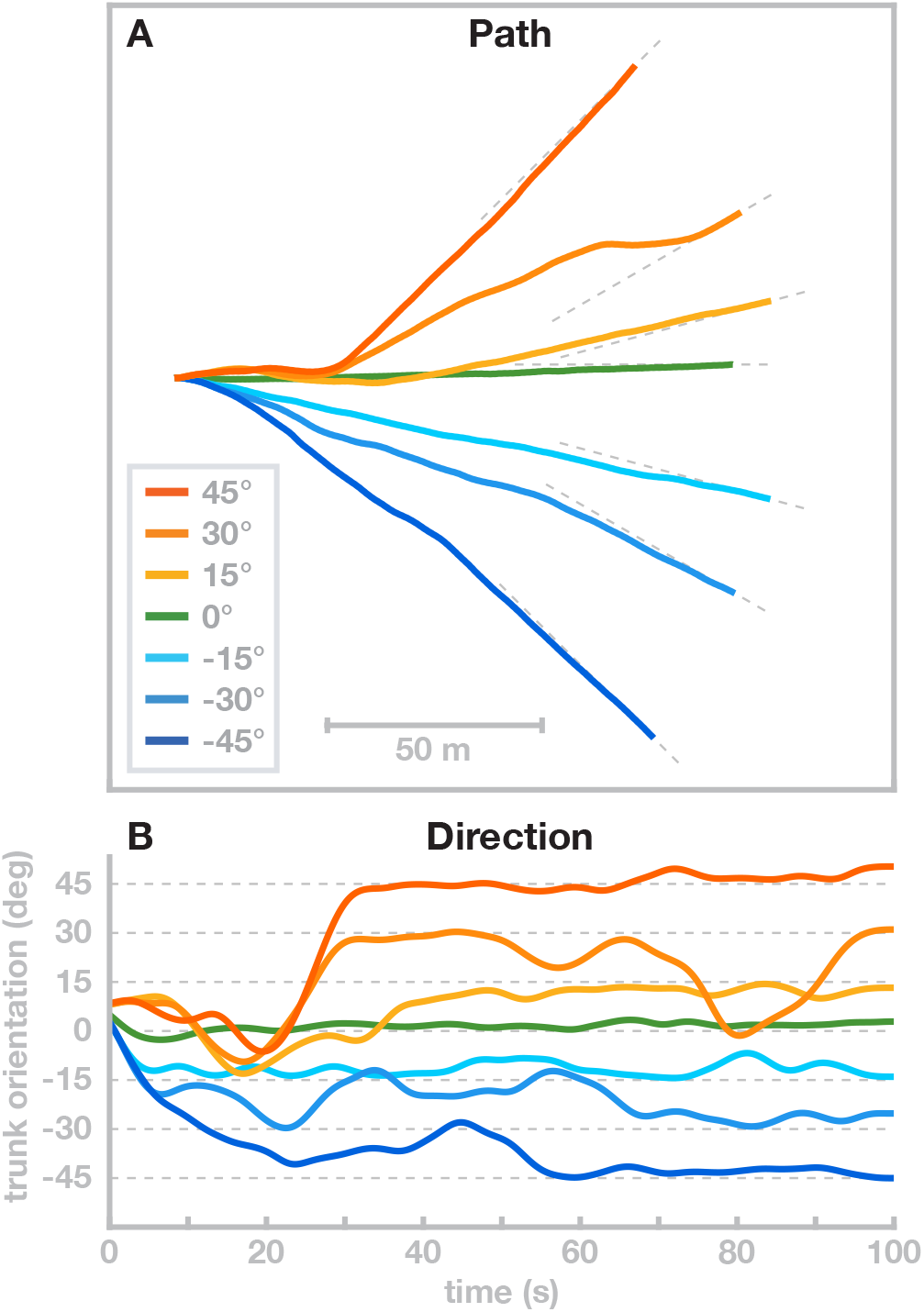
Direction control. Movement direction was adjusted by temporarily adding a constant offset value to the hip roll target angle until the target movement direction was reached, with seven different target directions. Panel A shows the horizontal path of the trunk center. Panel B shows the trunk orientation over time.

## 4 Discussion

We presented a musculoskeletal model of human locomotion that combines stable walking behaviour with the flexibility to generate voluntary movements with the swing leg according to a kinematic motor plan and to adapt the gait pattern. The model combines biomechanics, muscle physiology, spinal reflex loops and supraspinal neural processes in a physiologically plausible way. The supraspinal layer organizes the behavioural sequence, generates a movement plan on the task level and transforms the movement plan into descending motor commands that interface with the spinal cord. The spinal layer combines the descending motor commands for the swing leg with stretch reflex arcs for each muscle by shifting the muscle activation thresholds of the reflexes based on the descending command. Stance leg control is exclusively spinal, consisting of five dedicated reflex modules that each implement a specific function, following [105].

The model generates stable walking patterns, can flexibly move the swing leg according to a kinematic plan to avoid obstacles and can withstand external perturbations. Exploiting the passive properties of the model, we were furthermore able to change the model’s walking speed, cadence and movement direction to a limited degree.

### 4.1 Motor Plans and Voluntary Movements

The main innovation of the model we presented here is the ability to plan and execute voluntary movements with the swing leg, and to integrate these flexible swing leg movements into a stable gait cycle. Neuroscientists generally differentiate actions into two categories of *volitional* and *reflexive* actions [6]. Volitional actions are understood to be goal-directed, model-based and prospective, whereas reflexive actions are habitual, model-free, and retrospective [24]. Volitional actions are caused by a desire to reach a certain state in the future, whereas habitual or reflexive actions are caused by a stimulus in the past. While most of this work is at the intersection between neuroscience and psychology and investigates decisions, it intersects the field of motor planning and control.

Walking is largely considered a habitual movement, although it requires some executive control [18]. Decerebrated cats are able to walk without their brains, with only a tonic stimulation of their spinal cords [126]. Models of bipedal locomotion show that reflexes are sufficient in principle to generate stable walking patterns in principle [35, 31, 105]. The only high-level modulation required in these models is for balance control. A model of cat locomotion [130] shows that stable locomotion can already be achieved by a stereotyped rhythmic patterns from a CPG and state-dependent proprioceptive feedback is not critical, though it increases robustness of the gait pattern. These model results generally show that stable walking can be generated by low-level neural structures such as reflexes or CPGs. Precise, goal-directed movements, on the other hand, generally require cortical control. When receiving a motor cortex lesion, rodents and primates initially lose the ability to perform goal-directed reaching movements [127, 19]. Lesioned animals tend to recover some or large parts of the lost motor function over weeks or months after the lesion, either by local reorganization and neural plasticity [19] or by developing compensatory movements [32]. Even for goal-directed movement, the brain might not be critical. [51] showed that when rats learn a complex sequential lever press movement, they can still execute the learned movement after a lesion to the motor cortex. When receiving the motor cortex lesion before training, however, the rats were unable to perform or learn the lever press movement. Walking can be performed without requiring attention in steady-state on even or mildly uneven ground. More stringent constraints, such as walking over stepping stones or across a field cluttered with obstacles, require precise movements based on sensory information with a goal of getting the foot precisely onto a stepping stone, or around an obstacle [83, 15].

Existing models of walking are mostly reflex-based [35, 31, 105, 81, 23]. The walking movement pattern can be modified to some degree in various models to change speed or step over obstacles, but these modifications are designed for and limited to a specific target behavior. [112] shows shows that a walking model driven by a neural oscillator can adjust step length by adjusting timing and magnitude of the hip flexor activity, and increase toe clearance by superposing an additional descending motor command to the knee flexor muscles over the rhythmic activity. The model can step over obstacles placed at arbitrary positions by combining modulation of step length and toe clearance, but it lacks the control to move the foot along a specific path. [105] show that a model that is almost exclusively controlled by low-level reflexes can be generate stable walking movements in 3d. They achieve balance by modulating the reflex parameters slightly based on high-level information about the body in space. The model is robust in rugged terrain and has a certain degree of adaptability in that it can be made to walk at different speeds and change toe clearance to step over an obstacle. Adaptation is achieved by re-tuning the reflexes that map sensory information to muscle activation to a new cost function using evolutionary optimization [36]. Effectively, the model learns each behavior individually.

[120, 117] showed that it is possible to generalize between different sets of learned behaviors by interpolating between parameter sets, which generally results in an intermediate behavior. The mechanism can be used to combine the purely reflex-based walking generation in this class of model with a degree of central control, that maps a low-dimensional task parameter like walking speed onto a high-dimensional set of reflex parameters that will generate a walking pattern with the desired walking speed. But the movements generated by these models are still largely habitual, in that muscle activation is generated based on the current state of the system, rather than a desired future state and a motor plan for how to get to that state – they still lack the flexibility to plan and execute voluntary movements.

In the work presented here, we developed a model that combines reflex-based control of the stance leg with precise, goal-directed movements of the swing leg to generate walking movements that can be flexibly adapted to solve a task. Swing leg movements are planned on task level in the form of minimum jerk trajectories for kinematic task variables. The motor plan is represented as a trajectory that moves the task variable to a desired value in a specified time. For instance, swinging the leg forward is a planned movement of the thigh segment angle in the sagittal plane from a negative value at push-off to the positive value required for a successful heel-strike of the next step. This motor plan is updated during the movement to account for deviations from the planned trajectory of the task variable (see Section 2.1.4), and also to incorporate changes in the goal value required to maintain balance (see Section 2.1.3). To execute this motor plan, an inverse model of the spinal stretch reflex, muscle properties and biomechanics is used to calculate a descending motor command.

Our model has the flexibility to execute any movement plan as a volitional, goal-directed action. It can track random kinematic trajectories with high precision (see Section 3.1) when passively stabilized at the trunk. When moving freely, it can utilize this flexibility to move the leg over and around obstacles during swing. This flexibility is new for a walking model.

The model combines this flexibility with the ability to execute habitual movements, represented by sub-cortical reflexes that directly map sensory information to muscle activation. Stance leg control is completely reflex-based, while swing leg control combines the flexibility of goal-directed movements with the robustness of spinal stretch reflexes. The coordination of these two different types of behavior is organized by state-based switches. The neural mechanisms that implement this ability to smoothly swap be between different types of movement and sequentially combine habitual, reflexive control with volitional, goal-directed behavior are thought to be located in the basal ganglia [60]. Impairments to these structures, for instance from cell loss associated with Parkinson’s Disease, leads to reduced ability to switch between reflexive and goal-directed behavior, e.g. a reduced ability to voluntarily initiate gait from a standing posture, or the freezing of gait in some people with PD, which predominantly occurs in situations where environmental constraints require a goal-directed, planned modulation of a steady-state gait pattern, such as navigating through a doorway or over an obstacle [85, 124]. A mechanistic understanding of how impairments in neural function lead to specific motor deficits would require a model that encompasses both volitional and habitual movements, the neural mechanisms switching between them, and the integration with the spinal reflexes, muscle physiology and biomechanics that ultimately generate the movement. The model described here represents a first step towards such a mechanistic understanding.

### 4.2 Integration of High-level Control and Spinal Reflexes

For the swing leg control, our model uses a general stretch reflex that increases neural stimulation of the muscle based on the sensory information from muscle spindles about the length and velocity of the muscle [see Equation 16 and 62]. The descending command *u* shifts the set-point of this muscle-length feedback loop, and 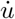 does the same for the velocity feedback. Similar equations have been used in various neuromechanical models of motor control, mostly of the upper limb [27, 35, 57, 13], but also in standing [95] and walking [35].

This control approach is a formalization of the general notion that reflexes are modulated by high-level input depending on task requirements. Studies in cats show that muscle spindle afferent activity is well predicted by muscle length and velocity [88]. This is consistent with the stretch reflex in Equation 16 with a fixed descending command. Other studies indicate that reflex gains can change depending on the phase of the gait cycle [134] and high-level tasks in both lower [68] and upper limb muscles [125]. While this evidence supports our assumption that spinal reflexes are modulated by supraspinal processes in task-dependent ways, the details of this interaction are not well understood experimentally and our control solution represents only one possible specification of this idea.

Technically, the formula we use is very similar to the equation used in the equilibrium point hypothesis approach to motor control [27]. This approach postulates that the spinal cord reflex modules simplify the control problem for the high-level areas, so that in order to move a limb to a desired position, the high-level controller only has to specify an equilibrium point corresponding to that position, and the low-level spinal reflexes generate the details of the actual movement [27, 13]. Modifications use different patterns of the descending command trajectory, like ramps or N-shapes [61, 33]. While more complex, these still adhere to the underlying concept that the structure of the descending command is simple and the spinal cord accounts for most of the complex details of the resulting movement pattern.

Despite the technical similarity in the stretch reflex, our model differs in the concept behind the equilibrium point hypothesis that the descending commands are simple. We found that considerable complexity is required to successfully generate movements that are both precise and flexible. One source of complexity are the highly non-linear inertial, gravitational and interaction forces that arise during locomotion. In a previous model of balance control in standing that with a similar control approach of shifting thresholds for stretch reflexes, we found that an internal model of the mass distribution and muscle moment arms across the joints and body segments was sufficient to maintain balance [95].

Internal models of body dynamics have widely been used in motor control theories to account for interaction forces [128, 52], often as components of optimal feedback control approaches [115], although the physiological plausibility of internal models with a complex dependency on the configuration state has been questioned [69]. Some approaches account for part of the biomechanics in an inverse model [89], and we follow this approach by modeling the swing leg dynamics, but not the compliant contact between foot and ground.

Still, the inertial forces alone are sufficiently complex to break the direct correspondence between task-level motor plan and muscle-level control, suggesting that an intermediate step is required to translate the high-level motor plan into descending commands.

The internal model used in our system to transform the high-level motor plan into low-level descending commands includes some aspects of the biomechanics, muscle properties and the stretch reflex. We do not claim that this specific solution is neurophysiologically plausible. Rather, we see it as a necessary connection between two systems with well-documented neurophysiological functions. There is good evidence that the higher motor areas in the brain plan and monitor movements using a task-level representation, e.g. the position or velocity of the hand when reaching to a target [30, 100, 16, 39]. There is similarly good evidence for low-level reflex arcs in the spinal cord, mapping proprioceptive signals directly to *α*-motorneuron activation [102, 53]. How the high-level movement plan is integrated with the low-level reflexes is currently not well understood [2, 4, 108].

In the present model, we used analytical inversion of the model equations and real-time re-planning for online updating to implement a module that functionally solves this problem of connecting the task-level motor plan with low-level motor areas in the spinal cord. We assume that this functionality is implemented neurally in the actual nervous system, solving the same problem but with a very different internal structure. There is some conceptual overlap with this notion and the equilibrium-point hypothesis, namely that there is a high-level motor control area that plans and generates movement on task level and then hands the details of execution over to more low-level structures. In walking, the present model shows, this transformation is of considerable complexity and needs to be addressed to generate movement patterns that actually walk. This likely points to a principled limitation of the equilibrium point hypothesis, that it does not account well for complex or time-dependent forces, as has been shown elsewhere [59, 38]. Interestingly, our model shows that this drawback of the equilibrium point hypothesis can be overcome by including internal models, two approaches that are usually seen as mutually exclusive.

### 4.3 Rhythmic Pattern Generation

In human walking, muscle forces, neural activity and ground reaction forces interact to generate rhythmic movement patterns. Existing approaches to model the dynamics of this combined system fall broadly in two categories, where the rhythmic neural pattern driving the motor system is either generated centrally [113], or emerges from the interaction between the body and the ground, fed back into the nervous system via sensory organs [105, 31, 81, 29, 122]. In the first approach, a dedicated neural structure, often called a central pattern generator (CPG), transforms a tonic neural activation into a rhythmic activation pattern between multiple neurons. CPGs are well-documented in insects [34, 70]. Evidence for CPGs has been found in cats, where a decerebrated cat can still walk when receiving tonic electrical stimulation at certain sites in the spinal cord [126]. [113] uses this approach to model human movement. In this model, a bank of neural oscillators drives the activation of the agonist-antagonist muscles spanning the leg joints, with one oscillator per joint. The structure of the neural oscillators broadly follows older models of spinal stepping generators [73, 50], consisting of two neurons, one activating the agonist and one the antagonist muscle of a joint. Such systems have stable oscillation patterns even in the absence of external inputs [71], though in Taga’s model both input and output are modulated depending on sensory data and the behavioral state, e.g. stance vs. swing. This model generates stable and robust walking patterns in the sagittal plane and can adapt to uneven terrain and additional loads [114]. Walking speed can be increased by adding tonic input and cadence can be controlled to a limited degree via entrainment by adding a rhythmic input.

In a second category of models, the rhythmic activity does not arise from neural oscillators, but from the interaction between neural control and the environment. In this class of models, muscle force generated by reflexes that drive a limb to a desired configuration, e.g. the swing leg forward after pushing off the ground. Different reflexes are turned on and off depending on sensory information, such as the leg switching from swing to stance once contact between the foot and the ground is detected. Organized appropriately, such interaction between reflexes, behavioral switches and environmental contacts generates stable oscillatory patterns. [119] showcase this principle in a biomechanically simple passive walker model with two legs actuated by spring-damper systems, where stable walking patterns emerge passively from the biomechanics, but the stiffness of the damped-spring muscles is increased at certain points in the cycle, based on sensory information, to replace the energy lost to damping back into the system. [35] use more realistic biomechanics with hip, knee and ankle joints that are actuated by Hill-type muscles, with muscle activation determined by generic stretch reflexes. Rhythmic patterns arise from switching between different set-points for the stretch reflexes, triggered by state feedback. Another model by [31] has similar biomechanics, but uses a selection of reflex modules to activate muscles. Each reflex module is designed to fulfill a specific function, activating a small set of muscles based on varied sensory input ranging from muscle length and velocity to forces and joint angles. [105] extended this model to 3d, and [120] combined it with a neural CPG.

The model presented here partially follows the tradition of combining reflexes with behavioral switches to generate rhythmic movement patterns. As some other models, our model shows a limited degree of flexibility in the generated walking patterns, where the resulting movement speed can be varied depending on the hip extensor force [114], the choice of control parameter set [105, 120], or in our case the trunk forward lean angle reference (see Section 3.3 above). [114] shows that cadence can be modulated as well by entraining the pattern generator to an external signal. The range in which cadence can be modulated in Taga’s model is relatively limited, spanning roughly 95–120 steps per minute. More recently, [23] showed that in a reflex model, modulation of a relatively small set of reflex parameters is sufficient to generate a wide range of walking patterns with cadences between 61–118 steps per minute, speeds between 0.48 and 1.71 m/s and step lengths between 0.43 and 0.88 m. While [114] varied cadence and speed together, [23] showed some independence, successfully modulating step length at a constant step duration, though failing to modulate step duration at a constant step length. When humans walk at a certain speed, they will generally use a certain combination of cadence and step length to achieve that speed that is largely invariant across repetitions [45]. But humans are also capable of walking at different combinations of cadence and step length for a given speed [80], as required e.g. when marching in-step. Our model can adjust speed to a limited degree by modulating the trunk lean angle and exploiting interactions between balance and speed (see Section 3.3). None of the currently existing models, our own included, is capable of this degree of flexibility. It can be argued that walking with a highly unusual combination of cadence and step length is more of a volitional action than normal walking, and requires motor planning and cortical control, which is largely absent in the existing models of human walking. This implies that our model should in principle be able to walk at any desired combination of cadence and step length, within a reasonable range, similar to humans. Achieving this would require a method to regulate speed by adding or removing energy from the system. Humans do this via push-off or hip extension of the stance leg [78]. To do the same, our model would need the ability to control the stance leg in the same goal-directed manner as the swing leg. While not possible in the model presented here, this is a promising direction for future research.

Our model does not contain a CPG. Forward movement of the swing leg is instead explicitly planned using a kinematic goal configuration and a minimum-jerk approach to plan a trajectory that transports the swing limb from the current state to the goal configuration. It is questionable whether humans generally use this high-level control to move the swing limb in a very deliberate, goal-directed way to a target during normal, steady state walking. We postulate that the human swing limb is controlled by spinal structures that can be modulated by supra-spinal inputs when necessary. In this model we implemented the spinal structures in the form of a general stretch reflex (Equation 16), which we then included in the internal model that transforms the kinematic movement plan into descending motor commands. It is an interesting question whether the stretch reflex in our model could be replaced with a CPG, which would be physiologically more plausible. While we see no reason why this should not be possible, the integration of the descending motor commands with a spinal CPG instead of a stretch reflex would be technically more challenging. This remains an interesting open question for future research.

### 4.4 Scope and Possible Extensions

We presented a neuromuscular model of human locomotion that combines flexible central control of the swing leg with fast and robust reflex-based control of the stance leg. Swing leg movements are realized as goal directed reaching movements and can easily adapt to required task constraints. Stance leg control, on the other hand, is achieved by five spinal reflex modules that (1) generate compliant, spring-like leg behavior, (2) prevent knee overextension, (3) balance the trunk, (4) compensate swing leg interactions and (5) plantarflex the ankle. This purely spinal control of the stance leg has the advantage that the leg can reactively compensate for unpredictable ground reaction forces on a fast time scale, without the need for central integration of different sensory systems, which is time consuming [84, 14, 118].

The presented model is limited such that the central controller has no direct access to the stance leg. Adaptations to desired stance leg motion patterns are only possible when reflex gain parameters are changed, requiring the re-optimization of the model parameters. Gaining high-level control over the stance leg could be achieved by superposing the existing reflex modules with additional descending control commands that realize desired gait adaptations while the functional reflex modules remain intact. The superposition of reflex modules and central control has been shown in a model of quite standing [111] where human sway signatures could be reproduced by combining muscle reflexes and virtual model control. We are currently working on extending the model in this direction to investigate if the superposition of descending and reflex-based control can be applied to the stance phase of locomotion.

Lateral balance control has been recently found to be governed by three biomechanical control mechanisms: The foot placement mechanism, the push-off-modulation and the ankle roll mechanism [91]. The foot placement mechanism describes an active shift of the lateral foot placement location at footfall after a perturbation [42, 12]. Shifting the footfall position changes the gravitational torque acting on the body through the new stance leg during the following step. This change in gravitational torque compensates for the perturbation. Push-off modulation is a change in the ankle flexion angle of the trailing leg during double stance, starting in late single stance [54, 55, 91]. An increase in the ankle plantarflexion, for instance, generates a push-off force that shifts the body weight between the two stance legs, in a direction that is largely forward, but also to the side [93]. The lateral component of the body weight shift compensates for lateral perturbations. The ankle roll mechanism is an active ankle inversion/eversion torque at the stance leg in single stance [41, 93], activating lateral ankle muscles to pull the foot segment and the rest of the body together. The foot segment rolls on the ground and shifts the CoP compensating for the perturbation. In the presented model, balance control solely relies on the foot-placement mechanism. This demonstrates that both push-off modulation and ankle roll mechanism are functionally not necessary for stable locomotion [116]. However, the two mechanisms are found to play a functional role in human walking increasing lateral stability especially in dedicated phases of the gait cycle. Simulations from simple SLIP models showed that using the ankle mechanism, when available, substantially reduces the amount of foot placement modulation required to maintain balance [92]. Adding the push off modulation and ankle roll mechanisms into the current model might improve balance in the model, leading to increased robustness against perturbations, and also lead to a better representation of human behavior by the model.

Human locomotion involves the coordination of multiple muscles spanning the different joints along the legs. Usually there are more muscles than biomechanical degrees of freedom, implying that there are different combinations of muscle forces that will lead to the same torques acting on the joints. Control requires selecting a particular solution out of this abundance of choice [9, 63, 103]. From a biomechanics perspective, specific muscles appear to be particularly appropriate for solving specific motor tasks. For instance, [40] showed that mono-articular muscles along the leg produce a force on the body center that is directed in the lengthwise direction along the limb, while the force from bi-articular muscles generates a significant transverse component. It is therefore biomechanically reasonable to compensate vertically acting gravitational forces with mono-articular muscles, while using bi-articular muscles when horizontal forces are required. E.g., the gastrocnemius muscle is mostly active during push-off, to propel the body mass forward, since this is one of the few situations where the combination of knee flexion and ankle plantarflexion generated by this muscle is functionally useful. Consistent with this general approach, neural evidence for the use of subgroups of muscles for balance control has been found by [99]. [99] showed that sagittal trunk stabilization during standing is mainly realized with biarticular hip muscles indicating that specific muscle groups might be dedicated to specific motor tasks. The use of muscle subgroup is generally considered as muscle synergies that have been found in walking [17, 47, 46] and reaching [20]. But how are these muscle synergies generated by the CNS? Spinal reflex circuits, as implemented in the stance leg in our model and several other models, map a sensory signal to a specific combination of muscles related to a functional motor task, e.g. stabilizing the knee. Even though multiple muscles affect one single joint, fixed reflex circuits define a unique combination of muscles that are recruited together. Such fixed reflex pathways, however, strongly restrict the ability of the limb to perform movements that are not captured by the pre-defined reflex, as discussed above. Specific co-activation patterns between muscles could also be realized by supra-spinal patterns, using specialized neural networks that learn an optimal solution to a specific task or sub-task that is encountered repeatedly with high frequency, such as swinging the leg forward during walking. In the present model we solved the mapping from joint torques to muscle forces in an ad-hoc manner using an iteration approach (see Section 2.1.5). Whether different solutions might provide functional benefits like improved stability or accuracy of voluntary movements requires further study.

## 5 Acknowledgements

RR and GS were funded by BMBF grant 01GQ1803. HR and JJJ were funded by the National Science Foundation grant CRCNS 1822568.

## Author contributions

Conceptualization: RR, HG, JJJ, GS, HR

Data Curation: RR, HR

Formal Analysis: RR, HR

Funding Acquisition: JJJ, GS, HR

Investigation: RR, HR

Methodology: RR, HG, GS, RR

Project Administration: JJJ, GS, HR

Resources: JJJ, GS

Software: RRR, HG, RR

Supervision: JJJ, GS, HR

Validation: RR, HR

Visualization: RR, HR

Writing – Original Draft Preparation: RR, HR

Writing – Review & Editing: RR, HG, JJJ, GS, HR

## Model sources

The model source files are available at https://github.com/hendrikreimann/FlexibleWalker.

## A Parameters

This sections lists the parameters used in the presented model. Table 1 shows the transport delays applied to the individual measures. The parameters used in the muscle-tendon and reflex model are listed in table 2. Table 3 shows the parameters used in the balance control equation and table 4 lists the parameters used for obstacle avoidance. The parameters of the ground-contact model are shown in table 5.

**Table 1:** Transport Delays

| Parameter | value (s) |
| --- | --- |
| $\delta_\theta$ | 0.01 |
| $\delta_{\dot{\theta}}$ | 0.01 |
| $\delta_{\ddot{\theta}}$ | 0.01 |
| $\delta_{\text{muscle length}}$ | 0.01 |
| $\delta_{\text{muscle velocity}}$ | 0.01 |
| $\delta_{\text{balance control}}$ | 0.1 |
| $\delta_u$ | 0.0025 |
| $\delta_{\dot{u}}$ | 0.0025 |
| $\delta_l$ | 0.0025 |
| $\delta_v$ | 0.0025 |

**Table 2:** Muscle and Reflex Parameters

| Parameter | value | unit |
| --- | --- | --- |
| $\tau_A$ | 0.01 | s |
| $h$ | 0.65 | - |
| $K_l$ | 5 | - |
| $K_v$ | 0.03 | - |

**Table 3:** Balance Control Parameters

| Parameter | value | unit |
| --- | --- | --- |
| $\phi_0$ | 0.44 | rad |
| $c_{d,\text{early}}$ | 0.47 | rad/m |
| $c_{d,\text{late}}$ | 0.30 | rad/m |
| $c_{v,\text{early}}$ | 0.17 | rad/(m/s) |
| $c_{v,\text{late}}$ | 0.2 | rad/(m/s) |
| $\phi_{0,\text{lat}}$ | 0 | rad |
| $c_{d,\text{early,lat}}$ | 0.13 | rad/m |
| $c_{d,\text{late,lat}}$ | 0.30 | rad/m |
| $c_{v,\text{early,lat}}$ | 0.31 | rad/(m/s) |
| $c_{v,\text{late,lat}}$ | 0.34 | rad/(m/s) |

**Table 4:** Obstacle Avoidance Parameters

| Parameter | value | unit |
| --- | --- | --- |
| $\alpha_{\text{hip}}$ | -1.5832 | rad/m |
| $\alpha_{\text{knee}}$ | -3.6941 | rad/m |
| $\alpha_{\text{ankle}}$ | 0 | rad/m |
| $\alpha_{\text{hiproll}}$ | -1.2 | rad/m |
| $c_{\text{hip}}$ | 2.3899 | rad |
| $c_{\text{knee}}$ | 2.3085 | rad |
| $c_{\text{ankle}}$ | 1.25 | rad |
| $c_{\text{hiproll}}$ | 0 | rad |

**Table 5:** Ground Contact Parameters

| Parameter | value | unit |
| --- | --- | --- |
| $\mu_{\text{static}}$ | 0.9 | - |
| $\mu_{\text{dynamic}}$ | 0.8 | - |
| Stiffness | 58860 | N/m |

